# Introducing divisive inhibition in the Wilson-Cowan model

**DOI:** 10.1101/2019.12.27.889642

**Authors:** Christoforos A. Papasavvas, Yujiang Wang

## Abstract

Both subtractive and divisive inhibition has been recorded in cortical circuits and recent findings suggest that different interneuronal populations are responsible for the different types of inhibition. This calls for the formulation and description of these inhibitory mechanisms in computational models of cortical networks. Neural mass and neural field models typically only feature subtractive inhibition. Here, we introduce how divisive inhibition can be incorporated in such models, using the Wilson-Cowan modelling formalism as an example. In addition, we show how the subtractive and divisive modulations can be combined. Including divisive inhibition in neural mass models is a crucial step in understanding its role in shaping oscillatory phenomena in cortical networks.

## 1 Introduction

Recent studies have demonstrated how different interneuronal populations deliver different inhibitory mechanisms in neocortex [1, 2]. Different inhibitory mechanisms differentially modulate the input-output function of the target neurons. For instance, subtractive inhibition hyperpolarise the target neuron by shifting the input-output curve to higher values of input, whereas divisive inhibition decreases the sensitivity (gain) of the target neuron by decreasing the slope of the input-output function [3, 4, 5]. In addition, divisive inhibition, as delivered by soma-targeting interneurons in neocortex [1, 2], decreases the maximum output of the target neuron along with the slope (see also, output divisive modulation in Discussion). Such modulations, among others effects, determine the dynamic repertoire of cortical networks and underlie neural computation [3].

Computational models can assist towards understanding the role of the different interneuronal populations and their inhibitory mechanisms in the dynamics of cortical networks. Abstract neural mass models, such as the Wilson-Cowan [6] and the Jansen-Ritt model [7], exhibit only subtractive inhibition [8, 9]. Introducing divisive inhibition in such models can help our understanding of its population level effect, and provide insights into its role in oscillatory phenomena in cortical networks.

Here we show how divisive inhibition can be introduced in the original Wilson-Cowan equations [6]. This is achieved by redefining the sigmoidal input-output function. In section 2, we formulate the input-output function that allows for a purely divisive inhibition and provide justification for its design. In section 3, we show how inhibition can be expressed as a combination of subtractive and divisive modulation with a more versatile input-output function.

## 2 Replacing subtractive with divisive inhibition

The Wilson-Cowan equations a set of two ordinary differential equations describing the interactions between an excitatory, *E*, and an inhibitory, *I*, neuronal population. They are expressed by

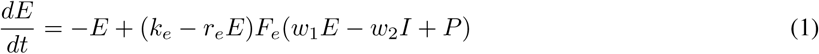

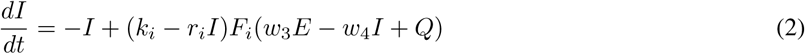

where *F*_*e,i*_(*x*), is a sigmoidal input-output function for the excitatory and inhibitory populations, respectively. This sigmoidal function introduces a non-linearity in each equation while its input is a linear combination of the weighted excitation, *wE*, the weighted inhibition, *wI*, and an external input, *P* or *Q*, to the population. The weights *w*_1,2,3,4_ are always considered non-negative. The equations are also equipped with a refractory period time constant *r*_*e,i*_ and the default maximum value of the input-output function *k*_*e,i*_ (see [6] for more details). Applying the logistic function as the sigmoidal function as suggested by Wilson and Cowan [6]:

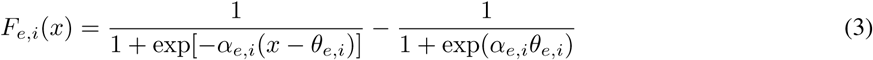

we get the following set of ODEs:

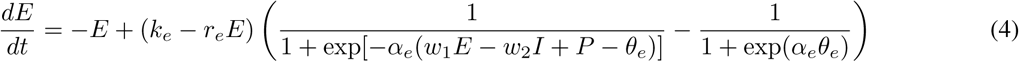

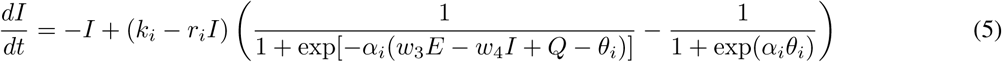

where the constant *α*_*e,i*_ > 0 governs the slope of the sigmoidal function whereas the constant *θ*_*e,i*_ > 0 represents the displacement of the curve on the x-axis (shift to the right). The sigmoidal function is shifted downward by the constant term 1*/*[1 + exp(*α*_*e,i*_*θ*_*e,i*_)] in order to achieve 0 output for 0 input, *F*_*e,i*_(0) = 0, following the formalism in [6].

Notice that the weighted activity level of the inhibitory population *wI*, which is always positive, is subtracted from the excitatory input. This is equivalent as shifting the input-output function to the right by increasing the parameter *θ*_*e,i*_ and this is why it should be considered a purely subtractive inhibition. Now, in order to transform it into purely divisive inhibition, we need to use it as a scaling factor of the sigmoid function instead (as in [10]). Thus, we substitute Eq. 4 with:

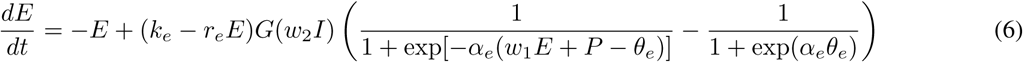

where function *G*(*x*) must be a monotonically decreasing function of *x*, where 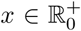, such that *G*(0) = 1 and it asymptotically converges to 0 for increasing values of *x*.

These requirements model the inhibition asserting a divisive effect, as the whole sigmoid is multiplied by a factor between zero and one, effectively decreasing the slope and maximum asymptotic value of the input-output function of the target *E* population. This modulatory effect of inhibition on the input-output function of pyramidal neurons was demonstrated in *in vivo* and *in vitro* studies [1, 2, 4, 5]. In such a setup, the magnitude of scaling should depend on the strength of the inhibitory population activity. Thus, an appropriate function for *G* would be:

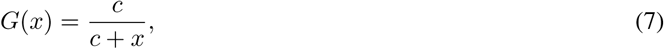

for any constant *c*. This constant can be set to the slope constant of the sigmoidal function *α*_*e,i*_, such that the sigmoidal is scaled down by a factor of 2 when the input of *G* reaches *α*_*e,i*_:

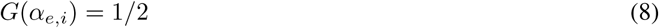

Thus the subtractive inhibition delivered to the excitatory population can be transformed into a divisive inhibition by replacing Eq. 4 with the following:

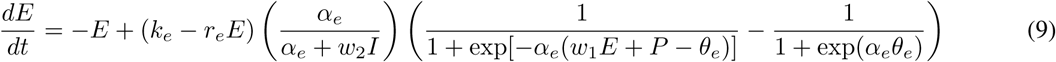

Similarly, the same transformation can be applied to the inhibitory population, if a divisive inhibition to the inhibitory population is modelled.

Equations 4 (subtractive inhibition onto *E*) and 9 (divisive inhibition onto *E*) can be reconciled by a generalised logistic function 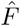 that has two additional inputs: a variable displacement Θ (subtractive modulator) and a variable scaling factor A (divisive modulator).

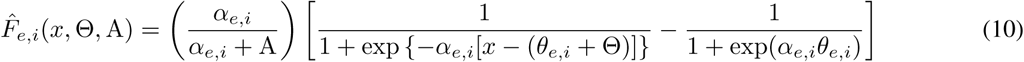

By using the generalised logistic function 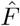, the following set of ODEs is equivalent to the original set in Eqs. 1-2 which considers inhibition to be purely subtractive

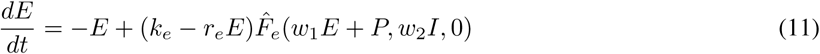

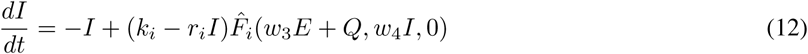

whereas the following set of ODEs considers inhibition to be purely divisive:

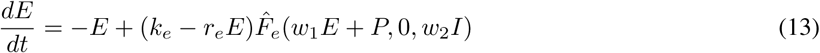

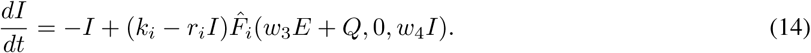

In this case, Eq. 13 is equivalent to Eq. 9.

## 3 Combining subtractive and divisive inhibition

An additional parameter can be introduced in the generalised logistic function 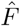 such that the divisive inhibition is no longer pure but, rather, a combination of divisive and subtractive modulation. Note this is conceptually and mathematically different to having two inhibitory populations delivering pure divisive and pure subtractive inhibition, respectively. This combination is important as a single inhibitory populations may not deliver purely subtractive or divisive inhibition, but rather a mixture.

By introducing an extra parameter 0 ≤ *q* ≤ 1 in the equation, it is possible to set the proportion of inhibition used as a divisive modulation, while the rest of it, 1 − *q*, is used as a subtractive one. This is expressed by:

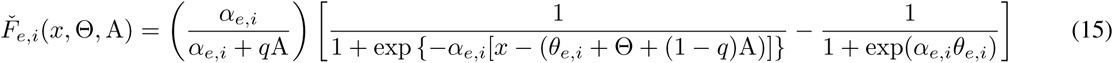

Notice that the subtractive modulator Θ is still purely subtractive but the divisive modulator A can range from purely subtractive for *q* = 0 to purely divisive for *q* = 1. This means that the original system in Eqs. 1-2 can be expressed by

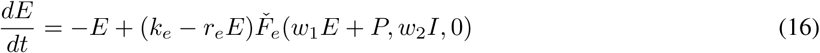

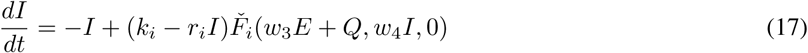

for any value of *q*. It can also be expressed by

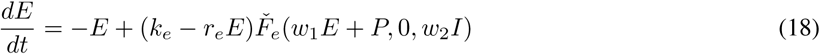

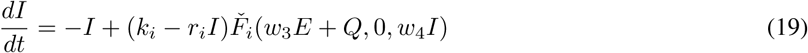

for *q* = 0. On the other extreme,

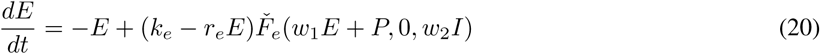

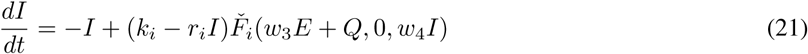

for *q* = 1, is equivalent to Eqs. 13-14.

As a side note, the redefined input-output functions above serve the same purpose but have a slightly different definition than the one used in [11].

## 4 Discussion

Above, we showed how divisive inhibition can be added to the original Wilson-Cowan model [6]. We also provide the justification for the design of a generalised version of the logistic function which allows for such an addition.

The generalised logistic function allows for the use of either subtractive or divisive modulation and, by using the divisiveness parameter *q*, it also allows for a mixed subtractive and divisive modulation. This means that Eqs. 18-19 can be used with varying values of *q* in order to create a continuum between the original system with purely subtractive inhibition and a system with purely divisive inhibition, with all the intermediate combinations of these two modulations, in order to investigate how divisive inhibition influences phenomena observed in the original Wilson-Cowan system. In addition, the generalised logistic function 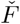 with the three inputs allows the description of cortical microcircuits with multiple inhibitory components with different modulatory effects (see soma-targeting vs. dendrite-targeting interneuron populations in neocortex [1, 2]). For example, one component can have purely subtractive effect and another component can provide a combination of subtractive and divisive effect, depending on the parameter *q* (for an example study, see [11]).

It is worth noting that what is described here is the output divisive modulation [3]. This means that the maximal output of the sigmoidal is decreasing along with the slope. A divisive modulation of the input would have an effect only on the slope. An alternative definition of the input-output function is needed for that (more similar to the one in [11] where the slope changes directly through the manipulation of the slope parameter of the logistic function). Here we focus on the output divisive modulation since this is the modulation that was reported to be delivered by the soma-targeting interneurons in neocortex [1, 2].

## References

[1] N. R. Wilson, C. A. Runyan, F. L. Wang, and M. Sur, “Division and subtraction by distinct cortical inhibitory networks in vivo,” Nature, vol. 488, pp. 343–348, aug 2012.

[2] B. V. Atallah, W. Bruns, M. Carandini, and M. Scanziani, “Parvalbumin-expressing interneurons linearly transform cortical responses to visual stimuli.,” Neuron, vol. 73, pp. 159–70, jan 2012.

[3] R. A. Silver, “Neuronal arithmetic,” Nat. Rev. Neurosci., vol. 11, pp. 474–89, jul 2010.

[4] M. Jadi, A. Polsky, J. Schiller, and B. W. Mel, “Location-dependent effects of inhibition on local spiking in pyramidal neuron dendrites,” PLoS Comput. Biol., vol. 8, no. 6, 2012.

[5] F. Pouille, O. Watkinson, M. Scanziani, and A. J. Trevelyan, “The contribution of synaptic location to inhibitory gain control in pyramidal cells,” Physiol. Rep., vol. 1, p. e00067, oct 2013.

[6] H. R. Wilson and J. D. Cowan, “Excitatory and inhibitory interactions in localized populations of model neurons.,” Biophys. J., vol. 12, pp. 1–24, jan 1972.

[7] B. H. Jansen and V. G. Rit, “Electroencephalogram and visual evoked potential generation in a mathematical model of coupled cortical columns,” Biol. Cybern., vol. 73, pp. 357–366, 1995.

[8] Y. Wang, M. Goodfellow, P. N. Taylor, and G. Baier, “Phase space approach for modeling of epileptic dynamics,” Phys. Rev. E, vol. 85, p. 061918, jun 2012.

[9] A. Spiegler, T. R. Knösche, K. Schwab, J. Haueisen, and F. M. Atay, “Modeling brain resonance phenomena using a neural mass model.,” PLoS Comput. Biol., vol. 7, p. e1002298, dec 2011.

[10] F. P. Battaglia and A. Treves, “Stable and rapid recurrent processing in realistic autoassociative memories.,” Neural Comput., vol. 10, pp. 431–50, feb 1998.

[11] C. A. Papasavvas, Y. Wang, A. J. Trevelyan, and M. Kaiser, “Gain control through divisive inhibition prevents abrupt transition to chaos in a neural mass model,” Phys. Rev. E, vol. 92, no. 3, p. 032723, 2015.

